# Alternative equations and “pseudo-statistical” approaches that enhance the precision of initial rates for the determination of kinetic parameters

**DOI:** 10.1101/2023.01.16.524223

**Authors:** Ikechukwu I. Udema

## Abstract

A burning concern among researchers studying enzyme kinetics has been ways of improving the accuracy of initial rates (*v*) with much greater precision. The goal of this study was to establish a formal (mathematical) way of achieving more accurate *v* values in enzyme assay. By adopting Bernfeld method of assay, the *v* values generated and other values in the literature were explored with two main objectives:1) to derive equations for a correctional calculation of initial rates and for the calculation of Michaelis-Menten (MM) parameters and 2) to evaluate the derived equations. The results of study showed that the maximum velocity (*V*_max_) and the MM constant (*K*_M_) obtained from a robust nonlinear regression (RNR) using *v* values in the literature were respectively between 21 and 26-fold and 6.56 to 61.2-fold greater than values generated from other method such as double reciprocal plot (drp), reciprocal variant of direct linear plot (RVDLP), and new method (z-model) derived based on MM equation; the *V*_max_ and *K*_M_ values (for alpha-amylase) calculated based on the RVDLP using the uncorrected *v* values were respectively 1.26- and 1.264-fold greater than the values using corrected *v* values; based on the *z* method, the *V*_max_ and *K*_M_ values using the uncorrected *v* values were 179.8- and 9.35-fold greater than the values using their corrected counterparts. In conclusion, the erroneous initial rates can be corrected using new equations before fitting RNR, RVDLP, drp, and any other equation to the corrected *v* values for the determination of MM parameters.

**Graphical Abstract figure:** 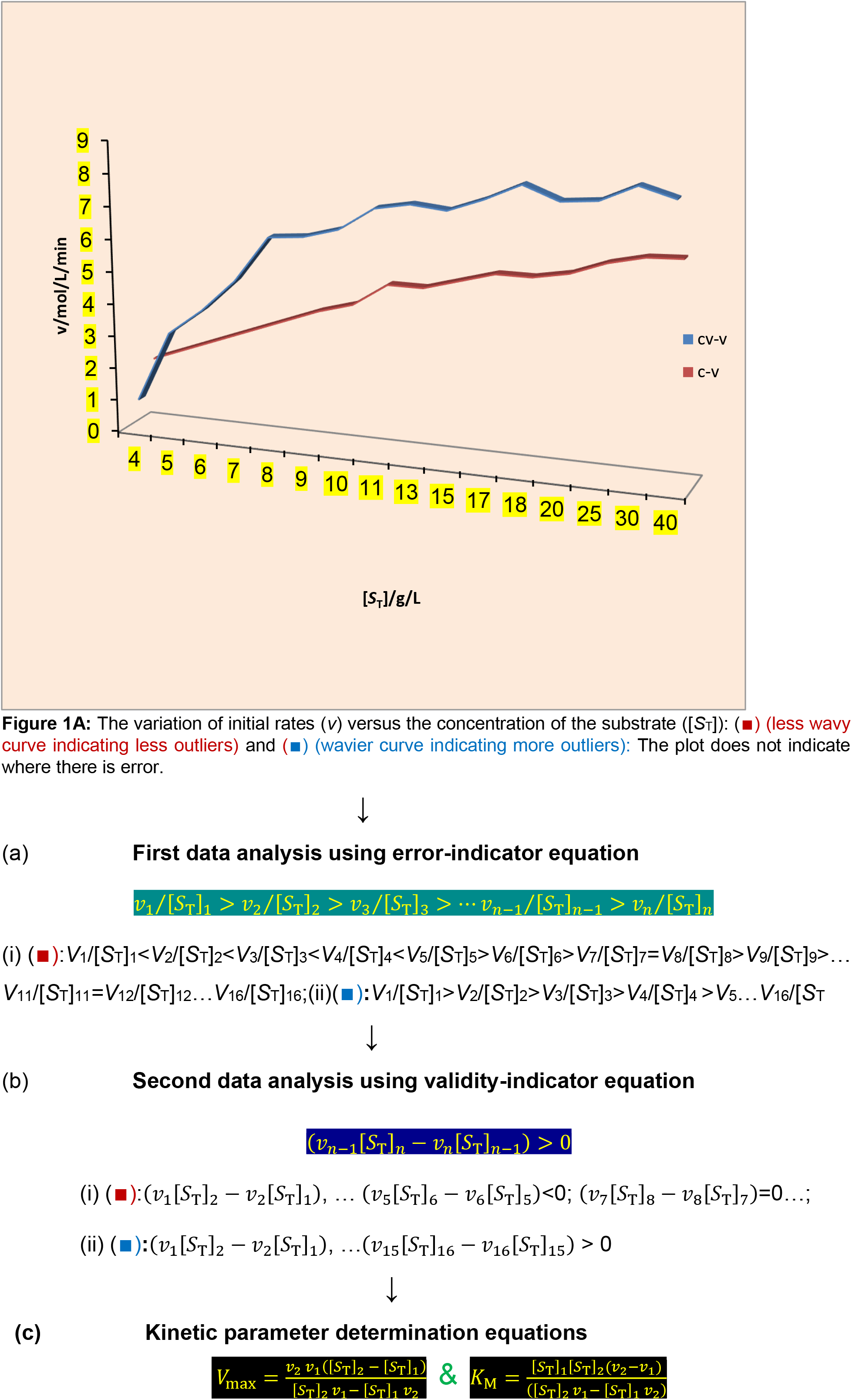

The implication of the result in b (i) is that the *V*_max_ and *K_M_* should either be a negative value or infinite value according to the equations unlike b (ii).

## Introduction

For a very long time, researchers have investigated ways in which the accuracy of experimental and kinetic parameters can be improved. The linearization of the original Michaelis-Menten [1] equation has been criticized because it yields inaccurate results due to outliers that are often ignored in most assays [2]. Inaccurate weighting of observations coupled with “outliers” at a higher frequency than allowed by normal distribution (perhaps Gaussian statistics) gives poor results [2]. It has been reported that the primary advantage of the Hanes-Woolf plot is that a plot of [*S*_T_]/*v* (where [*S*_T_] and *v* are the initial concentration of the substrate and rate of catalytic action) versus [*S*_T_] is not as heavily biased by velocities at very low [*S*_T_] as in the Lineweaver-Burk (LWB) plot [3]. The disadvantage of the Eadie-Hofstee (EH) equation is that the *v* appears on both sides of the equation; *v*/[*S*_T_] is not an independent variable [3]. But by the same line of argument, [*S*_T_]/*v* is not a dependent variable in the Hanes-Woolf (HW) equation. These equations are: 1/*v* = 1/*V*_max_ + *K*_M_*V*_max_/[*S*_T_] (LWB); [*S*_T_]/*v* = [*S*_T_]/*V*_max_ + *K*_M_/*V*_max_ (HW); and *v* = *V*_max_ – *K*_M_*V*_max_ /[*S*_T_] (EH). Matyska and kovář [4] investigated the most commonly used methods for estimating the parameters of nonlinear functions using simulated data with defined errors that have a structure similar to the errors of enzymological data obtained using standard techniques of initial-rate measurements. The authors strongly acknowledged that the Michaelis-Menten equation is a nonlinear equation and investigated various non-linear regression methods for fitting the Michaelis-Menten equation. With Marquardt’s method, the variability of the dependent variable (*i.e*., the initial reaction rates in our case) has to be taken into account. This variability is introduced into the calculation in the form of weights [5]. Given the square root of the variance (*σ*), the weight of each variable would have been 1/*σ* [5]. However, as in this study, only three data points are generated, which are not useful for Gaussian statistical analysis [4].

The introduction of the direct linear plot [6] and the suggested reciprocal variants [7] is a milestone in the desire to improve the accuracy of kinetic parameters. The existence of outliers is obviated by taking advantage of the nonparametric statistical approach, which involves counting rather than calculation for the estimation of the median value; however, the reciprocal variant requires minimal calculation. Thus, the direct graphical method can yield values similar to those obtained from computational procedures [6].

The motivation behind the desire to seek alternative ways of producing reliable Michaelian parameters by means of a robust nonlinear regression estimator based on a modified Tukey’s biweight function is the belief that outliers can lead to misleading values for the parameters of the nonlinear regression [8]. Similar beliefs and experiences have also prompted the suggestion for and use of software (MATLAB and any other software) for estimating enzyme kinetic parameters via the Wilkinson nonlinear regression technique [8, 9].

The goal of this study is to find a means of reducing the population of data points (pairs of *v* and [*S*_T_], for instance) to no more than two and yet achieve even more accurate estimates of primary kinetic parameters via alternative equations based on the Michaelis-Menten Equation and “pseudo-statistical” approaches that enhance the precision of kinetic parameters with the following objectives: The objectives of the study are: to 1) derive equations that are usable for a correctional calculation of initial rates if error is established; to 2) derive equations that define the incidence of experimental error in the initial rates; to 3) derive alternative equations, based on the Michaelian equation, for the calculation of the *K_M_* and *V_max_*; to 4) derive a pseudo-statistical equation for the minimization of the effect of experimental error; and to 5) evaluate the equation applicable to 1^st^, 2^nd^, 3^rd^, and 4^th^ objectives.

Let it be abundantly clear that either 6–15 (or more) replicates of each assay for different values of [*S*_T_] with a fixed concentration of the enzyme [*E*_T_] or 6–15 (or more) replicates of each assay for different values of [*E*_T_] can be done, but doing so may not yield results that are significantly different from duplicate studies or assays in the light of the new approach. In this study, the scope of work is scaled down to derivations of all equations and the determination of Michaelian parameters by calculation and graphical approach, to the exclusion of the pseudo-statistical approach for the correctional adjustment of parameters that have errors.

### Significance of study

Herein lies the significance of the study. Research is a very costly venture, even at a theoretical level, let alone at an experimental level where simulation may not be the case. A higher cost comes with the need for a larger data set. The need for precision and accuracy, in particular, cannot be sacrificed on the altar of cost-cutting. The new approach takes cognizance of this and can provide results with higher accuracy and validity at a lower cost. Every other method of generating Michaelian parameters, the direct linear plot and its reciprocal variants, and nonlinear regression approaches, do not correct the errors in the primary data, the initial velocities, but rather give an outcome based on the line of best fit, despite the fact that they represent a significant improvement in the accuracy and usability of the yielded kinetic parameters for whatever application they are used for, particularly the medical application. The new approach allows for the correction of the primary data and the initial rates based on the first principle inherent in Michaelian mathematical formalism, a nonlinear formalism.

## Theory

### Derivation of alternative forms of the equations for *v, K_M_*, and *V*_max_

Beginning with the known and progressing to the unknown in accordance with pedagogical principles, the first principle equation given by the two-source imputes of Michaelis-Menten and Briggs-Haldane takes into account the pieces of information in the literature [10]—the study of enzymes by Michaelis and Menten [1] by measuring *v* using Eq. (1a) below under the rapid-equilibrium assumption and the derivation of the equation by Briggs and Haldane [11] using the steady-state assumption.

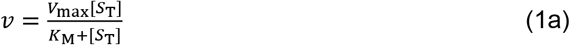

where *v*, *V*_max_, [*S*_T_], and *K*_M_ are the velocity of product formation (the initial rate), maximum velocity, concentration of substrate, and Michaelis-Menten constant, respectively. Another important view regarding the Michaelian equation takes into account the error function in the equation given as follows:

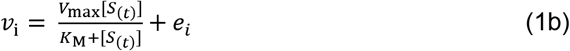

The *e*_i_ values are the random error components, and *n* denotes the sample size *(i.e*., the number of [*S*_T_], *v*_i_ pairs). *V*_max_ and *K*_M_ are the parameters whose values (and standard errors) are to be estimated; now, it is important to state that the authors recognize only [*S*_i_] (perhaps [*S*_T_] - [*S*_(t)_] with associated error) and *v*_i_ as steady-state parameters similar to observations elsewhere [12]. However, if errors are excluded from measured values of [*S*_T_], the only source of errors should be in *v*_i_, leading to errors in calculated or extrapolated values of *V*_max_ and *K*_M_. Without automated measurement of [*S*_T_], the errors in measured [*S*_T_] and *v* should translate into errors in *V*_max_ and *K*_M_.

Rearrangement of Michaelian equation (Eq. (1a)), given above, for two different velocities of enzymatic action (the initial rates) and the corresponding two different substrate concentrations results in the following.

First,

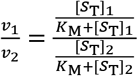

Rearrangement gives:

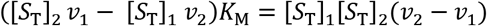

Then, making *K*_M_ subject of the formula gives:

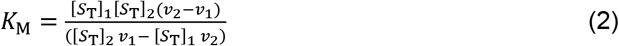

As stated in the text, there are shortcomings in the use of high-tech equipment in the measurement of the product of a reaction in most enzyme-catalyzed reactions. It is then obvious that common laboratory pipetting and timing with a stopwatch can be sources of error, resulting in a lot of outliers. To generate data (*i.e*. initial rates, *v*) with [*S*_T_], three different quantitative values of the substrate, S, can be substituted into the original Michaelis-Menten cum Briggs and Haldane equation, after generating the *K*_M_ and *V*_max_ by any known means, so that what should be the actual values of *v*, though theoretically determined, and corresponding to different concentrations of S, can be known. Note that, be it linear or non-linear regression or direct linear methods [2, 7], the generated *V*_max_ and *K*_M_ are not strictly equal to a linear functions of the velocities of enzymatic action; if they were, then a perfect condition should be the case. The line of best-fit actuated by either linear- or non-linear regression (and direct linear or its reciprocal variants) is purely a graduated improvement in the effect of the initial rate, *v*, on the generated kinetic parameters, *V*_max_ and *K*_M_, with direct linear (and its reciprocal variant) and non-linear regressions being superior to linear regression (the so-called linear transformations). Practical evidence is revealed in the result section. However, the first pre-cautionary measure is to run an assay using at least three different concentrations of the substrate for a Michaelian enzyme and substitute into an equation that gives, if not the exact value of *V*_max_, a close estimate; on the contrary, if such an operation is not possible *ab initio*, then there must be an error in the assay. The equation for inspection is given as follows: From the Michaelian equation, the derived equation is given as:

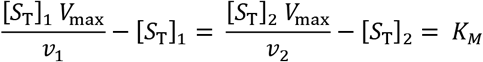

Rearrangement gives:

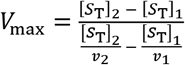

Simplification gives:

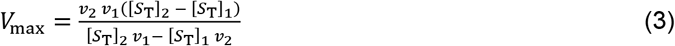

where *v*_1_, *v*_2_, [*S*_T_]_1_ and [*S*_T_]_2_ represent different velocities (*v*_2_>*v*_1_) of enzymatic action (amylolysis for instance) and different concentration ([*S*_T_]_2_ > [*S*_T_]_1_) of substrate. Effectiveness of Eq. (3) depends on the accuracy of the denominator which must be > zero or *v*_2_/[*S*_T_]_2_ must be accurately < *v*_1_/[*S*_T_]_1_. Equation (3) can accurately reproduce *V*_max_ if all measurements are made with very high precision, perhaps with automated instruments.

Rearrangement of the equation below [13] gives:

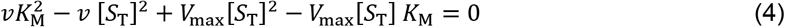

Division by *V*_max_ and further rearrangement gives:

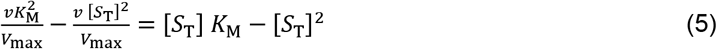

Equation (5) represents a general one. Now, consider two different velocities accurately corresponding to two different concentrations of S. One can now write for two different concentrations of the substrate as follows:

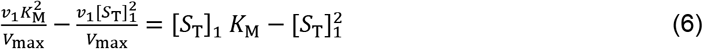

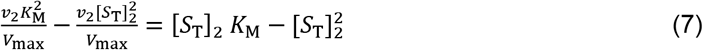

To eliminate 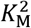, multiply Eq. (6) by *v*_2_ and Eq. (7) by *v*_1_. Solving simultaneously gives

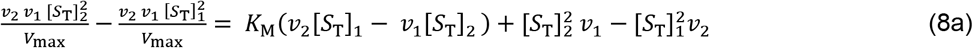

But, from Michaelian equation, one obtains the following: 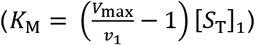. Note that [*S*_T_]_2_ and *v*_2_ can be used to give the same result. Substitute into Eq. (8a) to give:

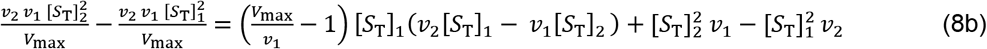

Rearrange and expand Eq. (8b) to give:

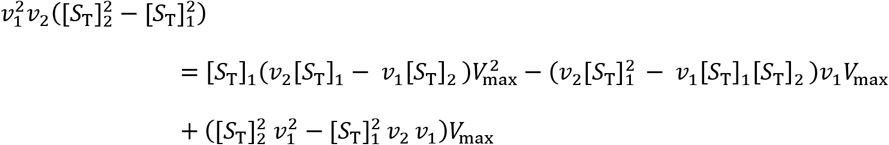

Rearrangement gives:

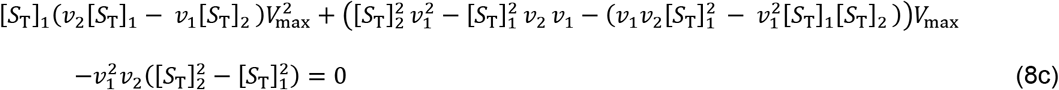

With change of sign for convenience, Eq. (8c) becomes:

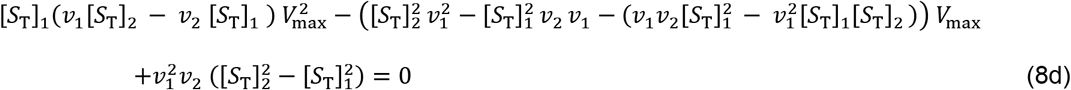

Since the root is likely to be too complex lets define the coefficients as follows:

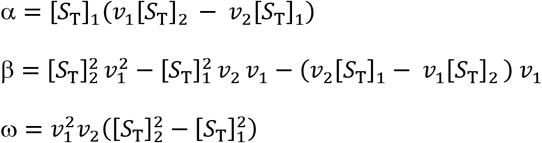

Then the solution is:

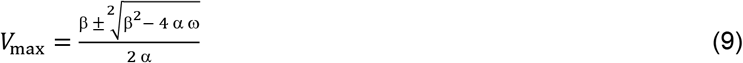

The positive root is validly and accurately a reproduction of the original *V*_max_ as long as *v*_1_/[*S*_T_]_1_ is accurately >*v*_2_ /[*S*_T_]_2_ or as long as *v*_1_ [*S*_T_]_2_ is correctly > *v*_2_ [*S*_T_]_1_.

From equation in the literature [13], is derived another form of the equation for the determination of *K*_M_ as follows: The equation in the literature is given as:

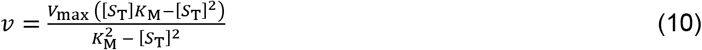

With Eq. (10) (or Eq. (8)), for different values of the velocity of catalysis, and with rearrangement, first via simultaneous equation approach), one gets

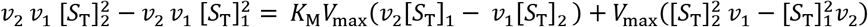

Defining *V*_max_ as 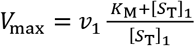 and substitution into the earlier equation leads to, after simplification and factorisation,

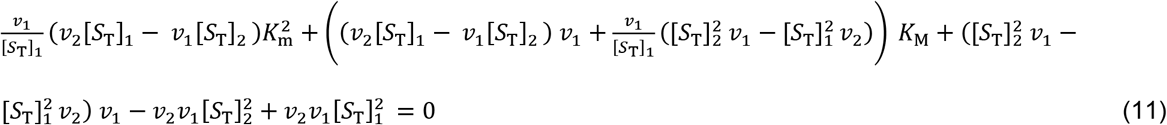

As in previous case, the roots may be too complex. Therefore, let

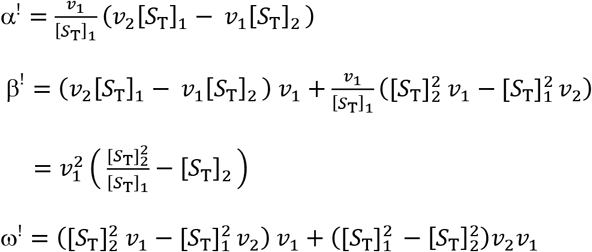

Such that,

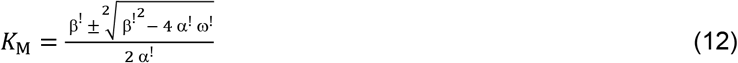

### Criteria for improved accuracy in the application of Michaelian kinetics and equation

The first criterion is the condition in which the following must hold:

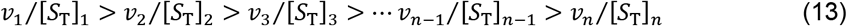

where, *n* is the number of pairs of (or single) assays carried out. The second criterion is the condition in which the following inequality must hold.

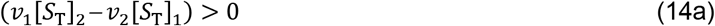

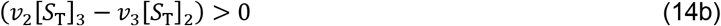

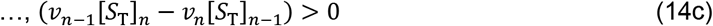

There is a need however, to point out that other forms of inequalities similar to Eqs (14a), (14b), and (14c) in terms of magnitude, are also applicable. Such is:

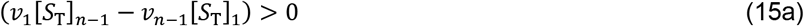

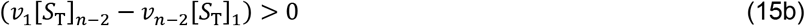

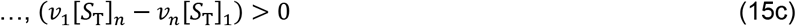

The third criterion is always as expected, even though the velocity of enzymatic action for a given S concentration, [*S*_T_]_n_, is < that for [*S*_T_]_n-1_, where both substrate concentrations are even > *K*_M_, thereby suggesting a possible product inhibition.

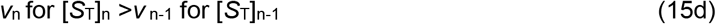

It must be noted that where the first criterion cannot be met, the second criterion cannot be met either, even if the third criterion is met. Where such an unfulfilled criterion is the case, a need for amendment arises, which can be achieved as follows: As a result, alternative equations to the Michaelis-Menten-Cum Briggs and Haldane equations, as well as pseudo-statistical approaches that aid in the precision of kinetic parameters, are required. In this regard, one can visualize the importance of the alternative equations, the first two, for instance, Eq. (2) for *K*_M_ and Eq. (3) for *V*_max_, as follows: If the denominator in both equations is less than one, equal to zero, or gives a negative value even if *v*_2_ is > *v*_1_, then the results generated cannot be completely error-free, regardless of the type of regression, linear or nonlinear, direct linear (or the reciprocal variant), *etc*. Thus, the general equation, Eq. (4c), is a better choice for ascertaining the absence or presence of very significant errors in the first three data points, *i.e., v*_1_, *v*_2_, and *v*_3_. If the three variables with their corresponding substrate concentrations are substituted into Eqs (2) and (3), the *K*_M_ and *V*_max_ may be very close to perfect values if, as previously stated, *v*_1_[*S*_T_]_2_ – *v*_2_[*S*_T_]_1_ is > 0; the same scenario is applicable to *v*_2_[*S*_T_]_3_ – *v*_3_[*S*_T_]_2_. If the variables were almost accurately determined by assay and substituted into the two equations, the values of the kinetic/Michaelian constant, if not equal, must be at least approximately equal. Every research project takes a significant amount of time and money to complete. But for the purpose of statistical analysis, a weighted data point [3] may be the beginning, but this demands replicate assays, up to a minimum of six (as a suggestion) for parametric statistical analysis; if ten different concentrations of the substrates are used, this would amount to sixty replicate assays.

Gaussian or parametric statisticians in the biological, paramedical, and medical sciences may prefer up to ten or more replicates, even if with just 4-6 replicates, a nonparametric approach may suffice. A mathematical model by Hozo *et al*., [14] is available for the determination of the standard deviation (SD). In fact, with 2-3 replicate assays for each substrate concentration ([*S*_T_]_1_–[*S*_T_]_3_), one can obtain 4-6 values of each of *K*_M_ and *V*_max_, just sufficient for a nonparametric analysis and the determination of the median and SD by Hozo *et al*. [14] method. The determination of the median for the estimates of *V*_max_ and *K*_M_ is also advised in the literature [15] in situations in which direct linear or reciprocal variant application cannot yield a definite or single intersection. It is thus worth noting that the *K*_M_ and *V*_max_ can be obtained with only three different [*S*_T_] and, as such, are expected to be very similar, if not identical, to what is expected from either a direct linear plot or nonlinear regression with a full range of substrate concentrations, ranging from 8 to 10 or more.

If *v*_1_[*S*_T_]_2_ – *v*_2_[*S*_T_]_1_ is not > 0 then the experimental variables must be re-calculated; as a precautionary measure, up to 4-5 experimental variables must be determined experimentally, but only in duplicate for the next two higher [*S*_T_]. There is a need for alternative equations when *v*_1_[S_T_]_2_ – *v*_2_[S_T_]_1_ is not > 0. The equations are not restricted to inaccurate *v*_1_, *v*_2_, *v*_3_, *v*_4_, *etc*. The first observation that has to be made is whether or not *v*_1_/[*S*_T_]_1_ is > *v*_2_/[*S*_T_]_2_; *v*_2_/[*S*_T_]_2_ is > *v*_3_/[*S*_T_]_3_, *etc*. If *v*_1_/[*S*_T_]_1_ is not > *v*_2_/[*S*_T_]_2_ but *v*_2_/[*S*_T_]_2_ is > *v*_3_/[*S*_T_]_3_, *v*_3_/[*S*_T_]_3_ is > *v*_4_/[*S*_T_]_4_, *etc*., then any of the latter can be used to correct *v*_1_ by calculation after substituting the more accurate *v* values and corresponding [*S*_T_] values into the derived equations. Such equations are derived as follows: Given any velocity of enzymatic action, the first corresponding to the lowest [*S*_T_], and any other velocity corresponding to a higher [*S*_T_], with the issue pertaining to the unfulfilled criterion in mind, the equation of maximum velocity to be determined by calculation in line with Eq.(3) is:

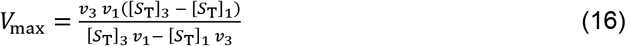

The appropriate equation of Michaelis-Menten constant, *K*_M_ in line with Eq. (2) is:

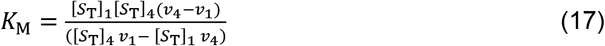

The equation for *K*_M_ is chosen as presented because it is derivable from velocity, *v*_4_, which is higher than *v*_3_, which is considered to have higher precision. Equations (16) and (17) can then be substituted into the Michaelis-Menten equation for *v*_3_ to give:

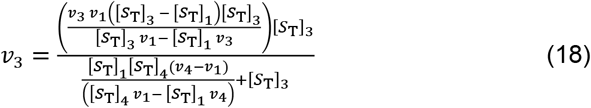

Following tortuous rearrangement one gets

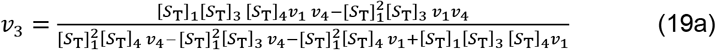

Eq. (19a) simplifies to:

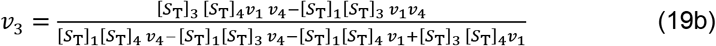

From Eq. (19a), one gets the equation for *v*_1_ which may have problem of lower precision. Again following tortuous rearrangement one gets:

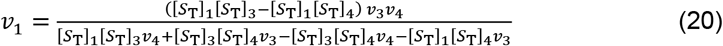

Equation (20) may over estimate the value of *v*_1_. Therefore, values of [*S*_T_] and *v*, that are chosen, are those immediately lower than those that appear in Eq. (20) to yield:

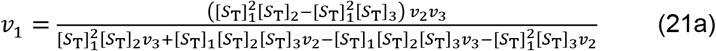

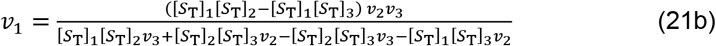

If error in *v*_2_ is the case the equation that can apply is derived as follows:

First, the *K*_M_ equation is:

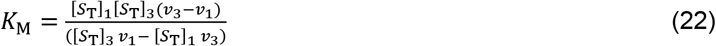

And the *V*_max_ equation is:

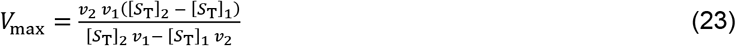

The same operations that led to Eqs (18) and (19), applied to Eq. (19), leads to:

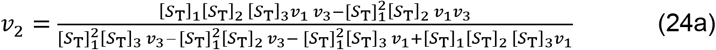

Eq. (24a) simplifies to:

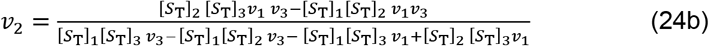

Any other higher velocity with imprecision can be treated intuitively in the same way, with the *caveat* that the immediate higher velocity to be chosen is the one after the lower velocity where there is error that often leads to outliers. In other words, the next higher velocity after the velocity (ies) where there was an error must be substituted into Eqs. (2) and (3). Equations (9 and 10) may be avoided because of their complexity and implied time consumption index. Some of other equations are derived as follows: The equation for *v*_2_ is considered first. The equation for *K*_M_ is one in which *v*_2_ should not appear, such that:

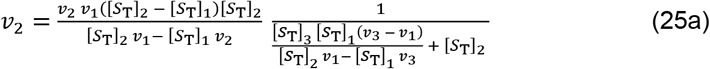

Expansion gives:

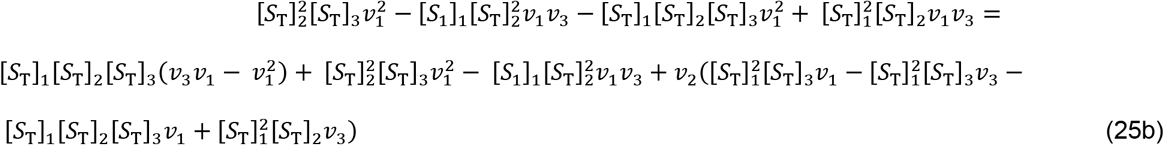

Cancellation of common factors and rearrangement gives:

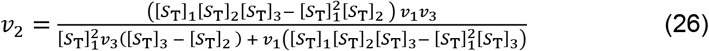

Rearrangement of Eq. (26) or by going through the same procedure leading to Eq. (26), one gets:

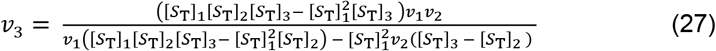

In applying Eq. (27), the recalculated values of *v*_2_ and *v*_1_ may be substituted into it for the calculation of *v*_3_. This rule is applicable to any other case(s).

The equations for *v*_4_, *v*_5_, and *v*_6_ can be derived following the same argument and procedure such that:

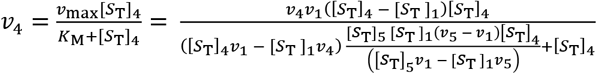

Simplifying gives:

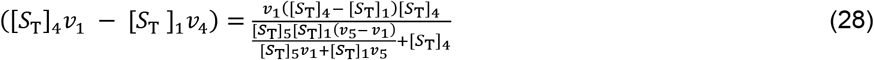

Expanding Eq. (28) and making *v*_4_ subject of the formula gives:

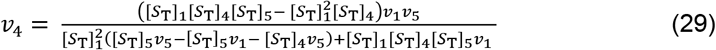

Equation for *v*_5_ is derived by making it subject of the formula in Eq. (29).

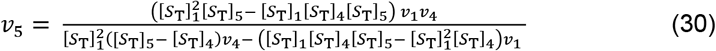

Equation for *v*_6_ can be derived directly from Eq. (30) by changing measured and experimental variables.

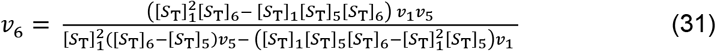

Any results based on Eqs. (2) and (3), respectively, for *K*_M_ and *V*_max_ that are much higher than results obtained from both linear transformation and direct linear, nonlinear, *etc*. methods demand that the raw data be subjected to pseudo-statistical treatment. A quasi-weighting treatment needs to be applied to the data according to the following equations, derived according to the method of trial and error until very suitable equations were obtained. For clarity and contrast, *K*_M_ and *V*_max_ are rewritten as 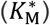 and 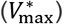, which stand for imprecise (error-laden) *K*_m_ and *V*_max_, respectively.

Thus,

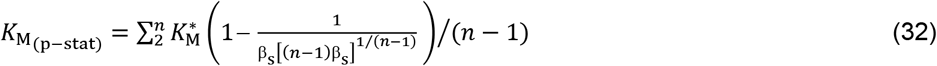

where, *K*_M(p-stat)_, and *n* are the mathematically and pseudo-statistically determined *K*_M_ and the maximum number of different [*S*_T_]respectively. The coefficient, β_s_ is taken to be a weighting factor for the fractional contribution of each substrate to the excess concentration observed in the summation result.

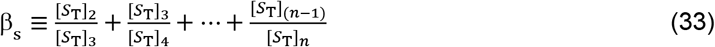

The outcome of assay is first and foremost, the product formed; hence a second weighting factor (**β_p_**), the product type, is given as:

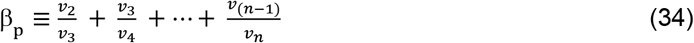

The summation 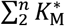 is given in either of two ways such as:

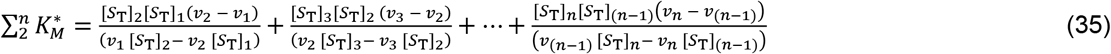

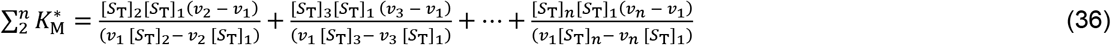

Equations (26) and (27) can give the same result if the assay is performed without error, particularly where high-tech automated equipment is used for the assay. But in cases of error-laden results, different results are likely, and they may be abnormally high. The mathematical and pseudo-statistical equations for determining *V*_max_, restated as *V*_max(p-stat)_, are as follows:

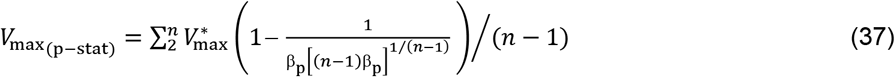

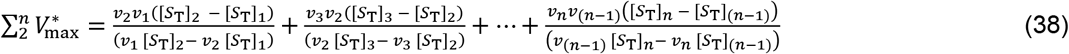

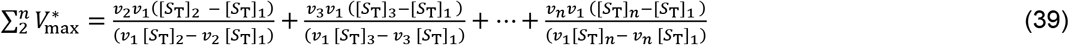

Equations 38 and 39 can give a similar result if the experimental variables contain minimal or negligible error in each variable. Perfect data cannot easily be generated in routine laboratory assays, whether academic, clinical, diagnostic, or otherwise.

Equations (14a) through (15c) must hold in order that Eqs (25) and (28) can be applied. If the *V*_max_ is calculated using a double reciprocal plot (drp) which is known [3] to overestimate the kinetic parameters when the raw data is error-laden, the following pseudo-statistical weighting technique may be used.

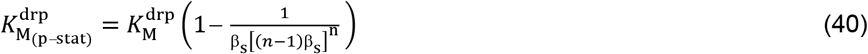

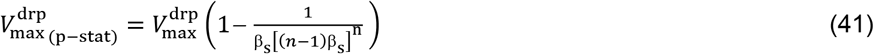

With respect to Eqs (40) and (41), the weighting factors take, respectively, the form:

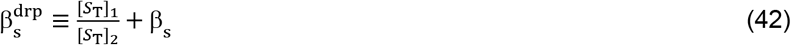

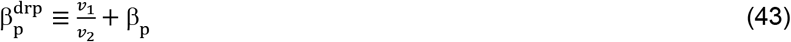

This means that β_S_ and β_P_ in Eqs (40) and (41), respectively, have to be replaced with Eqs (42) and (43) where the Michaelian parameters are obtained by double reciprocal plots.

It needs to be made clear that none of the equations for the recalculation of the initial rates of catalysis should be held sacrosanct. Rather, one needs to subject them to rational ratiocination. The first step is the application of Eqs (13) and (14c); if the criteria are met, then one should calculate the values of either the *V*_max_ or the *K*_M_ for different pairs of values of [*S*_T_], namely [*S*T]_1_/[*S*_T_]_2_, [*S*_T_]_1_/[*S*_T_]_2_, [*S*_T_]_1_/[*S*_T_]_2_;…[*S*_T_]_(n-1)_/[*S*_T_]_n_. The calculation can still be carried out even if the criteria are not met; the action enables the identification of points where there are errors for correction. From the calculated values of *V*_max_ and *K*_M_, one may identify values of either *V*_max_ or *K*_M_ that are significantly larger than the average; the values of the *v* that produced such a larger value have to be calculationally corrected by substituting the *V*_max_ and *K_M_* values that are lower than the average into the correctional equations. Such equations may not necessarily be two or more of the equations derived above; a little patience and intuition are all that are needed in deriving suitable correctional equations.

## Results

To begin with, let it be known that subjecting rates to linear regression and nonlinear regression analysis gives results that are more indicative of the line of best fit than a direct linear plot or analysis. However, the most important issue is the need to realize that the original rates (the so-called initial rates, as a matter of clarity) cannot be the same as the values of the same parameter obtained by calculation if the *V*_max_ and the *K*_M_ are substituted into the first principle Michaelian equation. These recalculated rates are exactly the values that can reproduce the *V*_max_ and *K*_M_ values by whatever regression method. This is to imply that if there are three sets of data, each should give different values of the Michaelian parameters; substitution of each of the pair of parameters, *K*_M_ and *V*_max_, into the Michaelian equation gives, after calculation, different values of the rate per different [*S*_T_]. This is important for the calculation of other secondary but vital kinetic parameters usually obtained by calculation or by the graphical method [16, 17].

As illustrated in Figures 1, 2, 3, and 4, Eqs (1) and (2) are actually nonlinear equations in that the observed values of the coefficient of determination (R^2^) resulting from the linear plot with the “uncorrected” (unweighted) *v* values (Figure 1 and 2) are < the value for the polynomial plot (Figures 3 and 4), which is suggestive of nonlinearity in the relationship between *v* and [*S*_T_]. The plots (Figures 1 and 2) with corrected initial rates compares in a similar fashion-the values of R^2^ for the linear plot were < the values for the polynomial plot (Figures 3 and 4) as applicable to the literature data and in this study. However, there is a slight deviation with respect to alpha-amylase in this study in which the R^2^ values for the linear (Figure 2) and polynomial (Figure 4s) plots with the unweighted (“uncorrected”), *v* values are equal.

**Figure 1:**
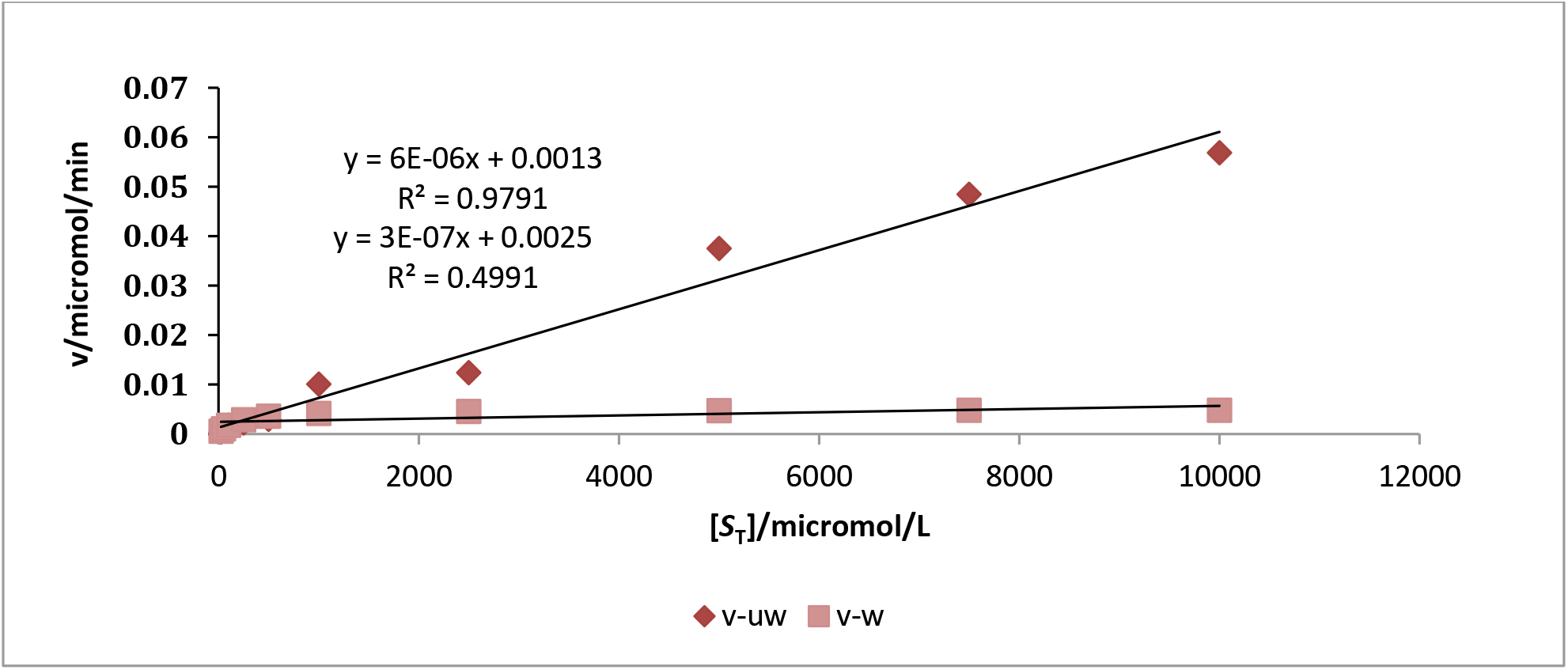
A linear plot of initial rates (*v*) [8] versus [*S*_T_] (different concentrations of L-DOPA) verus [*S*_T_] for the evaluation of linearity characteristic of pre-steady steady and steady-state far from zero-order kinetics: (■) denotes the plot for the “weighted” (or the corrected) *v* versus [*S*_T_] and (**◆**) denotes the plot for the un-weighted *v* versus [*S*_T_].

**Figure 2:**
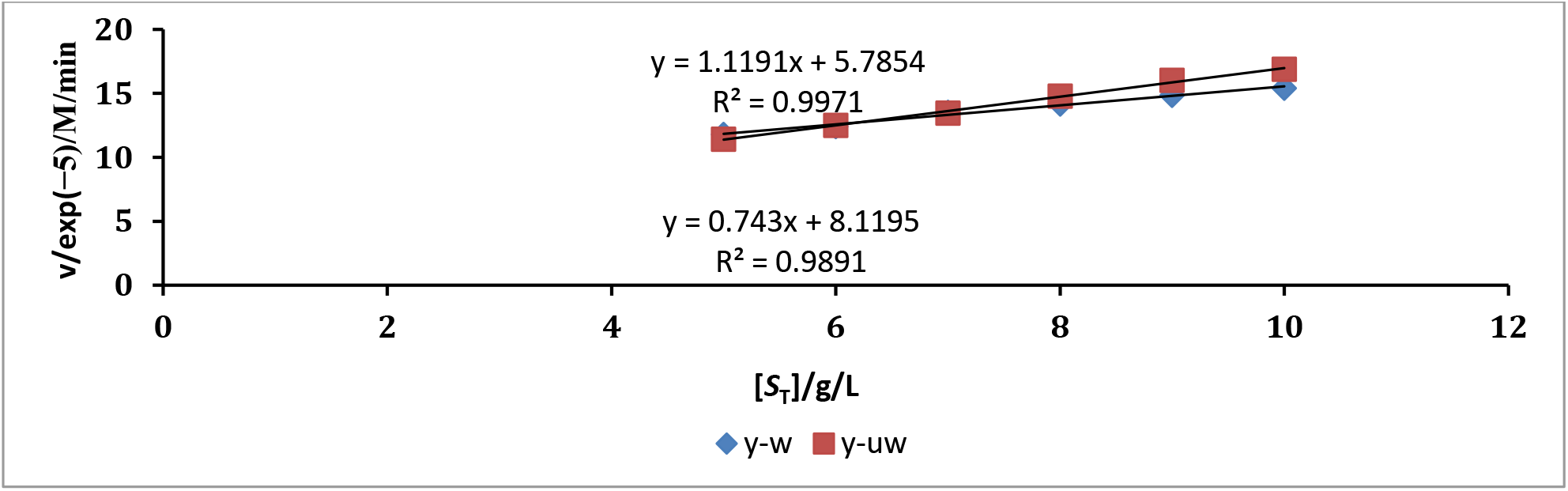
A linear plot of initial rates (*v*) in this study versus [*S*_T_] (different concentrations of gelatinized in soluble potato starch) verus [*S*_T_] for the evaluation of linearity characteristic of pre-steady steady and steadystate far from zero-order kinetics: (**◆**) denotes the plot for the “weighted” (or the corrected) *v* versus [*S*_T_] and (■) denotes the plot for the un-weighted *v* versus [*S*_T_]. The enzyme assayed is alpha-amylase.

**Figure 3:**
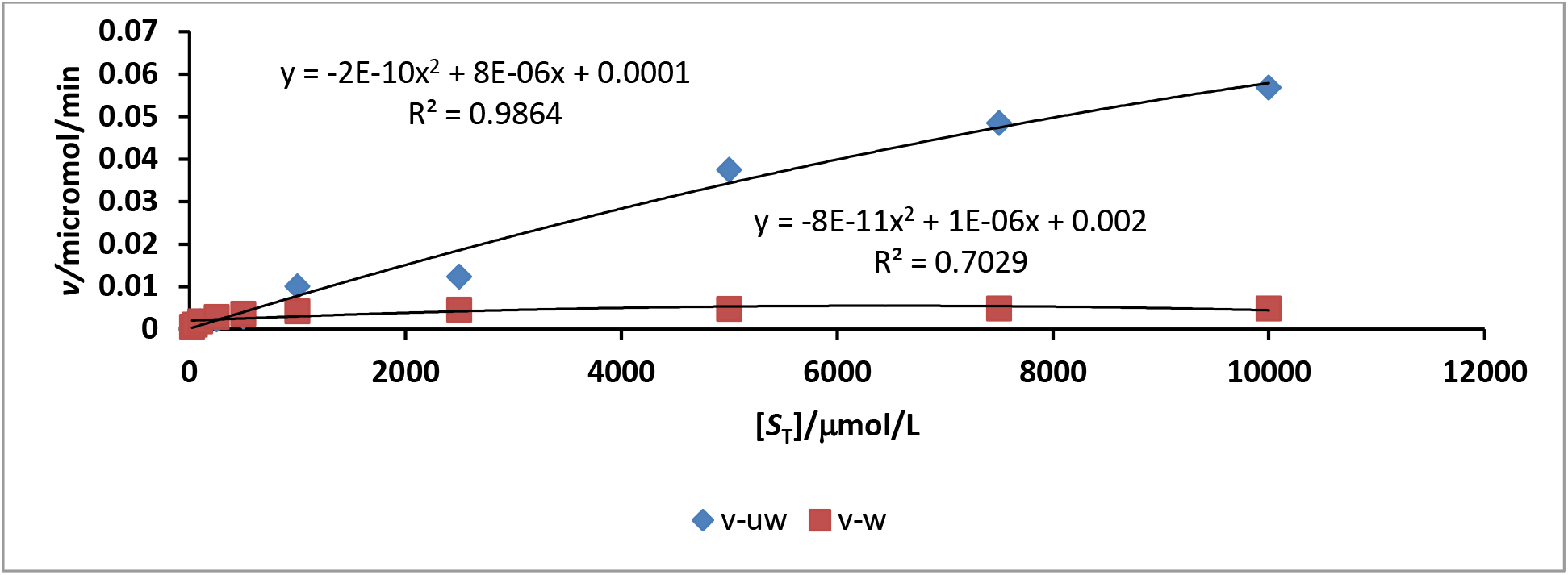
A polynomial plot of initial rates (*v*) [8] versus [*S*_T_] (different concentrations of L-DOPA): (■) denotes the plot for the “weighted” (or the corrected) *v* versus [*S*_T_] and (**◆**) denotes the plot for the un-weighted *v* versus [*S*_T_].

**Figure 4:**
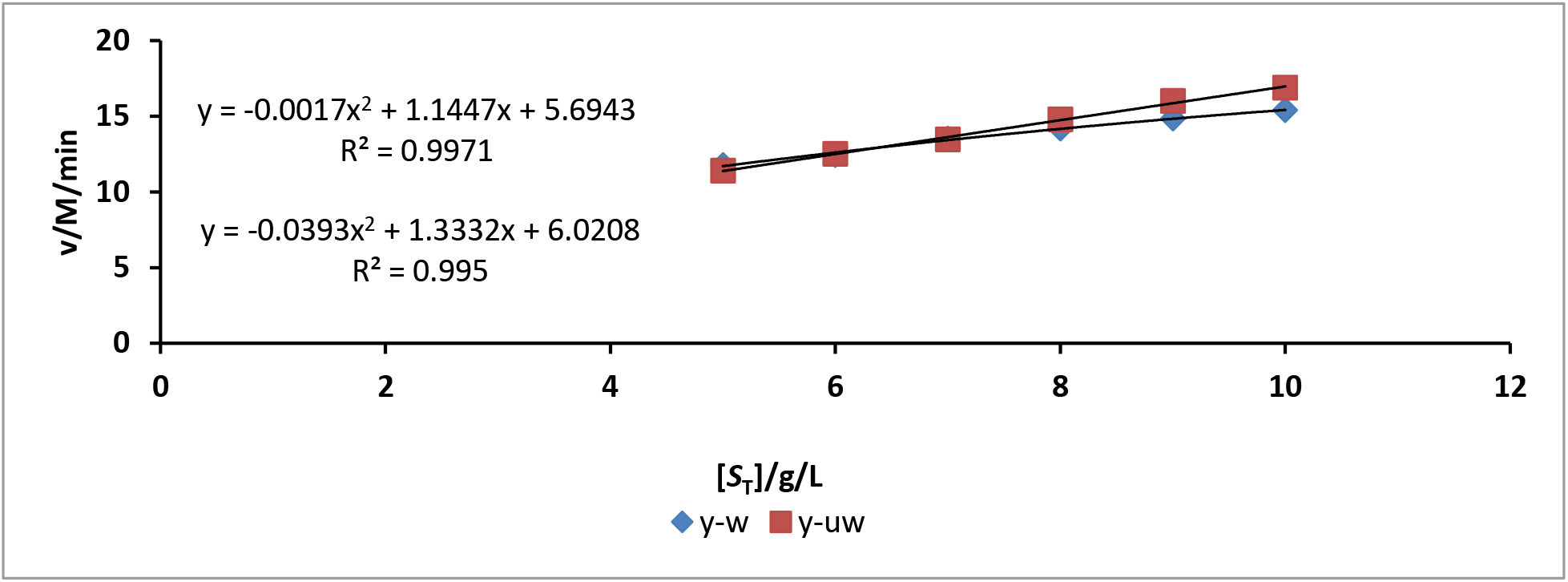
A polynomial plot of initial rates (*v*) in this study versus [*S*_T_] (different concentrations of gelatinized in soluble potato starch) verus [*S*_T_] for the evaluation of linearity characteristic of pre-steady steady and steady-state far from zero-order kinetics: (**◆**) denotes the plot for the “weighted” (or the corrected) *v* versus [*S*_T_] and (■) denotes the plot for the un-weighted *v* versus [*S*_T_]. The enzyme assayed is alpha-amylase.

The temptation on the part of any one is to feel that the plots so far are too elementary in the pursuit of much higher postdoctoral research, particularly by those neck deep in the mathematical sciences; one should not be surprised to observe that flawed research reports may be occasioned by the negligence of the fundamentals in the area of research. For instance, if catalytic rate and its reverse rate counterpart were determined under the conditions that validate reverse quasi-steady-state approximation (rQSSA) and were found to be very efficient and effective in the activation of drugs *in vitro* for possible application *in vivo* using an animal model, any future report based on the Michaelian model or the standard QSSA (sQSSA) may be flawed on two grounds. Because the higher concentration of *in vivo* enzymes is known to have an *in vitro* counterpart whose much lower concentration is as predetermined by the experimenter, administering drugs or any other medicinal formulation to experimental animals can produce effects that are greater than expected, with adverse effects in some cases, if a similar concentration of drug as in an *in vitro* scenario is administered. Data generated from this type of scenario can be considered invalid or flawed. A guide based on sQSSA in the formulation of drugs, solid and liquid, ignores the fact that the *in vivo* concentration of the target enzyme is *»* open-ended choice of different concentrations *in vitro*, though most often any of such concentrations prepared is ≪ *in vivo* concentration of the same enzyme. A simple overall massage is that, the foundation of accurate kinetic parameters is an accurate measurement of initial rates coupled with the correct information about the steady-state assumption that best defines the kinetic parameters.

For the purpose of a further comparison, one may look at the leading terms of the linear and polynomial curves; the slope of the linear curve for the unweighted *v* values, the outliers notwithstanding, is steeper than for the corrected values (Figure 1). It goes to show, *ab initio*, that there should have been a trend towards Michaelian kinetics if not for the outliers due to experimental errors in *v* values. The plots (Figure 2) for alpha-amylase showed similar differences. A steeper slope is still an expression of pre-steady-state and steady-state kinetics behind the zero-order limit. On the other hand, the coefficient of the leading terms in the polynomial plot or equation is numerically higher with unweighted *v* values than with the corrected *v* values (Figures 3 and 4); this is applicable to Tyrosinase (EC1.14.18.1)[8] and alpha amylase. Generally speaking, one needs to realize that a plot of *v* versus [*S*_T_] does not correctly represent the Michaelis-Menten equation; rather, it is more likely to represent the equation: *v* = *V*_max_ [*S*_T_]/*K*_d_, where *K*_d_ is the dissociation constant of the enzyme-substrate (ES) complex. Most standard text books and journals retain *K*_M_ rather than *K*_d_.

Nonetheless, the linear and polynomial plots with the corrected (“weighted”) *v* values yielded different R^2^ values, the polynomial plot giving the higher value. The initial rates in the literature [8] were used to execute the graphical illustration as shown in Figures 1 and 3 while Figures 2 and 4 were created in this study in which *Aspergillus oryzae* alpha amylase is a test model. The literature data was chosen because it contains a wider range of substrate concentrations and was subjected to a robust nonlinear regression for the estimation of kinetic parameters. The data obtained can be used for the evaluation of the results of this study in a comparative manner. Meanwhile, the reciprocal variant of the direct linear plot is explored for the generation of the Michaelian parameters that are regarded as a reference standard for comparative assessment of the values of the parameters obtained from a double reciprocal plot and the robust nonlinear regression analysis of the data from the literature [8] only. Of course, a polynomial plot as opposed to the software-orchestrated nonlinear regression is not a better approach. Its use in this study is an illustration only.

Figures 3 and 4 reaffirm the inappropriateness of a linear plot in that the use of weighted or corrected *v* values resulted in a poorer correlation with a linear plot compared to polynomial plot even if as stated, polynomial plot is of lower value than nonlinear regression. Deductively, one can opine that, the more the data assumes a Michaelian characteristic, otherwise known as a hyperbolic relationship between *v* and [*S*_T_], the linear and polynomial plots become less relevant and less appropriate with poorer correlation. Of course, the excellent correlation between *v* and [*S*_T_] in a linear regression analysis is simply an expression of kinetics far from zero-order kinetics, where Michaelian kinetics is relevant. One needs to add that perfect linearity is an absolute indicator of the realm of kinetics that “justifies the reverse quasi-steady-state approximation and vice versa”. The graphs reveal that the ratio of [*E*_T_] to [*S*_T_] may not be ≪ 1.

At this juncture, in the light of Eqs (1a) and (1b), it is necessary to point out the fact that whenever linear regression and nonlinear regression are carried out, they do not take any leverage on those equations because in those forms, the initial velocities are simply fractions of the maximum velocities. A plot of *v* versus [*S*_T_] does not take into account the denominator, which contains another [*S*_T_]. This is where the reciprocal variant of the direct linear plot becomes very useful in that it takes into account two variables stated as a ratio such as [*S*_T_]/*v* in which one is a function of the other (higher concentration of [*S*_T_] translates into higher concentration of the product with time (Table 1 and 2); the reservation against this method may include the suggestion that [*S*_T_] is not an independent variable [3]. However, there is a need to realize that with the mass conservation principle in mind, the remaining substrate concentration is a function of the initial rates, which in turn are a function of the efficiency of the biological catalysis to act hydrolytically on the substrate in accordance with the nature of the latter. This implies that the remaining substrate concentration in time *“t”* (the duration of the assay) is some sort of dependent variable. Hence, a direct linear plot (reciprocal variant) of 1/*v* versus ([*S*_T_] – *t v M_p_*)/*v* can address the concern in the literature. This discussion is important because the reciprocal variant of the direct linear plot was explored for the generation of kinetic parameters that are serving as a standard reference for the evaluation of similar parameters in this study and in the literature. Nevertheless, there is also the concern in the literature [18.] that a direct linear plot where the number (*n*) of data points, such as the pairs of *v* and [*S*_T_], is > 5 is impracticable; this has prompted the desire for software that can address the challenge. Nevertheless, improvization cannot be ruled out in research; this was the case in this study, in which the reciprocal variant of the direct linear plot was executed with the aid of Microsoft Excel.

**Table 1:** Values with and without corrections that verify the accuracy or otherwise of the initial rates from the literature [8] for the Tyrosinase-catalyzed reaction.

|  |  |  |  |  |  |  |  |  |  |  |
| --- | --- | --- | --- | --- | --- | --- | --- | --- | --- | --- |
| $\ddagger^{(c)}v/[S_T]$<br>(e. (-6)) | 24 | 22 | 18 | 12 | 7 | 5 | 4 | 1 | 0.6 | 0.5 |
| $\ddagger^{(uc)}v/[S_T]$<br>(e. (-6) mol./g) | •16 | 22 | 18 | 9 | 6 | •10 | 5 | •8 | 6 | •6 |
| $\ddagger^{(c)}\Delta\vert v[S_T]\vert /e. (-15)$<br>(mol<sup>2</sup>/L<sup>2</sup>.min) | 5.94 | 65 | 284 | 539 | 1561 | 5906 | 11174 | 11958 | 48326 | - |
| $\ddagger^{(uc)}\Delta\vert v[S_T]\vert /e. (-15)$<br>(mol<sup>2</sup>/L<sup>2</sup>. min) | -7.5 | 20 | 22 | 400 | 1500 | 12850 | -31750 | 38750 | 58250 | - |
| $\ddagger^c v$<br>(nmol/L) | 0.62 | 1.80 | 2.30 | 2.91 | 3.67 | 4.21 | 4.63 | 4.78 | 4.84 | 4.87 |
| $\ddagger^{uc} v$<br>(nmol/L) | 0.40 | 1.10 | 1.80 | 2.30 | 3.00 | 10.10 | 12.40 | 37.50 | 48.50 | 56.90 |
| % $e_{rr}$<br>e. (+2) | 0.4 | 0.4 | 0.22 | 0.2 | 1.8 | -1.4 | -1.7 | -7 | -9 | -11 |

The percentage error is expressed as the difference between the corrected and uncorrected rates divided by the corrected initial rate, exp. (+2). The ash tag, as superscript indicates, data from the literature [8] "err" and "e." denote error and exponent, respectively; ‡ denotes literature values of the initial rates. $\Delta|v[S_T]| = ([S_T]_n v_{n-1} - v_n [S_T]_{n-1})$ ; c and uc denote corrected and uncorrected, respectively. \* indicates the presence of error in the initial rate

**Table 2:** Values with and without corrections that verify the accuracy or otherwise of the initial rates in this study for the alpha-amylase-catalyzed reaction.

|  |  |  |  |  |  |  |
| --- | --- | --- | --- | --- | --- | --- |
| $^{(c)}v/[S_T]$<br>(e. (-5)) | 2.28 | 2.09 | 1.92 | 1.77 | 1.65 | 1.54 |
| $\ddagger^{(uc)}v/[S_T]$<br>(e. (-5) mol./g) | 2.28 | 2.09 | 1.92 | 1.85 | 1.79 | 1.67 |
| $\ddagger^{(c)}\Delta\vert v[S_T]\vert /e. (-5)$<br>(mol<sup>2</sup>/L<sup>2</sup>.min) | 5.74 | 7.22 | 3.98 | 8.98 | 9.75 | - |
| $\ddagger^{(uc)}\Delta v[S_T]/e. (-5)$<br>(mol<sup>2</sup>/L<sup>2</sup>. min) | 5.74 | 7.22 | 3.98 | 4.18 | 10.38 | - |
| $\ddagger^c v$<br>(μmol./L) | 114 | 125 | 134 | 142 | 148 | 154 |
| $\pm_{uc} v$<br>($\mu\text{mol./L}$) | 114 | 125 | 134 | 148 | 161 | 167 |
| % $e_{rr}$<br>e. (+2) | 0.0 | 0.0 | 0.0 | -0.042 | -0.088 | -0.084 |

The percentage error is expressed as the difference between the corrected and uncorrected rates divided by the corrected initial rate, exp. (+2). "err" and "e." denote error and exponent, respectively. $\Delta|v[S_T]| = ([S_T]_n v_{n-1} - v_n [S_T]_{n-1})$ ; c and uc denote corrected and uncorrected, respectively.

The main concern in this study is the issue of accuracy, not necessarily perfection, though striving to achieve perfection should always be the overriding goal, even if it is never attainable. As Table 1 shows, the initial rates reported in the literature have errors marked in black superscript as evaluated using Eq. (13). There was an irregular trend in the values with respect to increasing substrate concentrations (25–10,000 mmol./L); this clearly violates the rule of accuracy (and possibly validity) as expressed in Eq. (13). On the contrary, the corrected version complied with the rule of accuracy. It is important to state categorically that neither the direct linear plot (nor its reciprocal variant) nor the nonlinear regression can correct the errors; rather, they produce results for Michaelian parameters that are closer to the real values, if, in particular, the weighting scheme applied in the nonlinear regression is accurate. The test of the validity or accuracy of the difference in the denominator (Eq. (15c)), showed mixed results, containing negative and positive values and irregularity in trend with respect to increasing substrate concentration. This is, as expected, a clear violation of the rule of accuracy applicable to the “uncorrected” (unweighted) initial rates as opposed to the corrected version which occasioned a regular trend (Table 1). This should form the initial background check before embarking on a full-scale assay with multiple replicates for therapeutic applications. One can correctly deduce that increasing trend of initial rates with increasing concentration of the substrate is not always without error as evidenced by the percentage errors in Table 1.

Preliminary observation (Table 2) showed that *v*_1_, *v*_2_, and *v*_3_ were relatively accurate; *v*_4_ and *v*_7_ were not very accurate. Although *v*_3_/[*S*_T_]_3_, *v*_9_/[*S*_T_]_9_, and *v*_10_/[S]_10_ demonstrated a normal decreasing trend, this is insufficient to ensure that the expression of difference (Eq. (15c)) is consistently positive and in the correct ratio with the nominator (*v*_n_*v*_n–1_ ([*S*]_n_ – [*S*]_n-1_) in order to give a consistent value of *V*_max_ and consistently, *K*_M_. It appears, therefore, that the observed decreasing trend defined by Eq. (13) is not a conclusive indicator of accurate values of the Michaelian parameters that may be calculated or graphically determined. However, the first two tests of accuracy and possibly validity have been fulfilled. Yet they need to be tested when applied to the calculation of the Michaelian parameters.

The general forms of Eqs (2) and (3), [*S*_T_]_n_[*S*_T_]_n–1_(*v*_n_-*v*_n-1_)/([*S*_T_]_n_*v*_n–1_-*v*_n_[*S*_T_]_n-1_), and *v*_n-1_*v*_n_([*S*_T_]_n_-[*S*_T_]_n–1_)/([*S*_T_]_n_*v*_n-1_-*v*_n_[*S*_T_]_n–1_) can be explored for the calculation of the *K*_M_ and *V*_max_, respectively. As Table 3 shows, the values of the Michaelian parameters obtained by fitting relevant equations to the raw (“uncorrected,” otherwise referred to as “unweighted”) data are *»* than the values calculated by exploring the corrected initial rates. The values of the *V*_max_ and *K*_M_ from exploring the corrected values of *v* are 4.95 nmol./L/min and 175 μmol./L, respectively, while the corresponding values from exploring unweighted *v* values are 25.1 nmol./L/min and 1636 μmol./L; these values are ≪the corresponding values, 208.6 nmol./L/min and 25720 μmol./L obtained from ordinary least square regression (OLS) [8] as well as 118 nmol/min and 10740 μmol./L obtained from robust nonlinear regression (RNR) [8]. A confirmatory double reciprocal plot gave results (*K*_M_ = 1201 μmol. /L and *V*_max_ = 20.79 nmol. /L/min) very similar to the literature report [8]; both results showed values that were again ≪ values obtained from OLS and RNR. The reference standard values (i.e. values derived from the reciprocal variant of the direct linear plot (RVDLP) which are *V*_max_ = 4.7 nmol./L/min and *K*_M_ = 208.3 μmol./L which are values from exploring corrected *v* values and the corresponding values such as 5.26 nmol./L/min and 263.16 μmol./L obtained by exploring uncorrected *v* values are even exceedingly ≪ the values obtained from OLS and RNR in the literature [8].

**Table 3:**
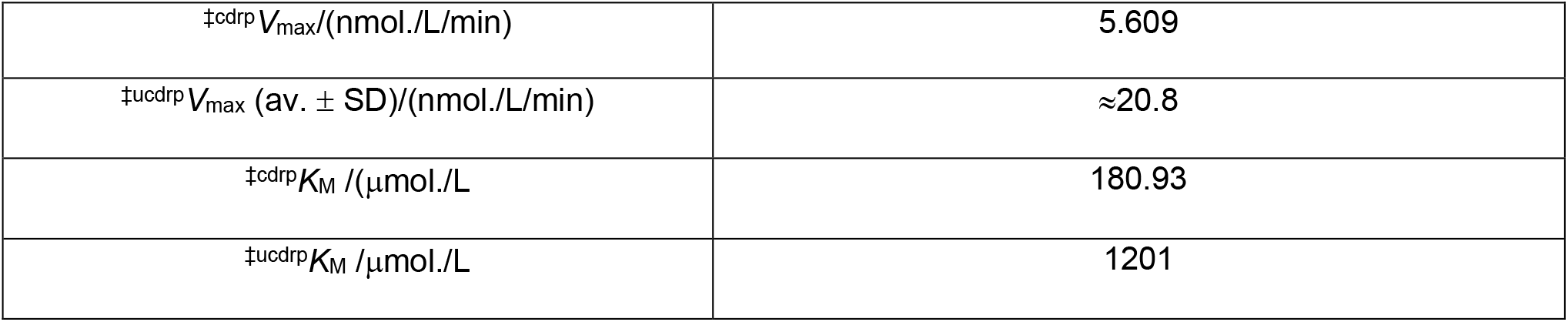
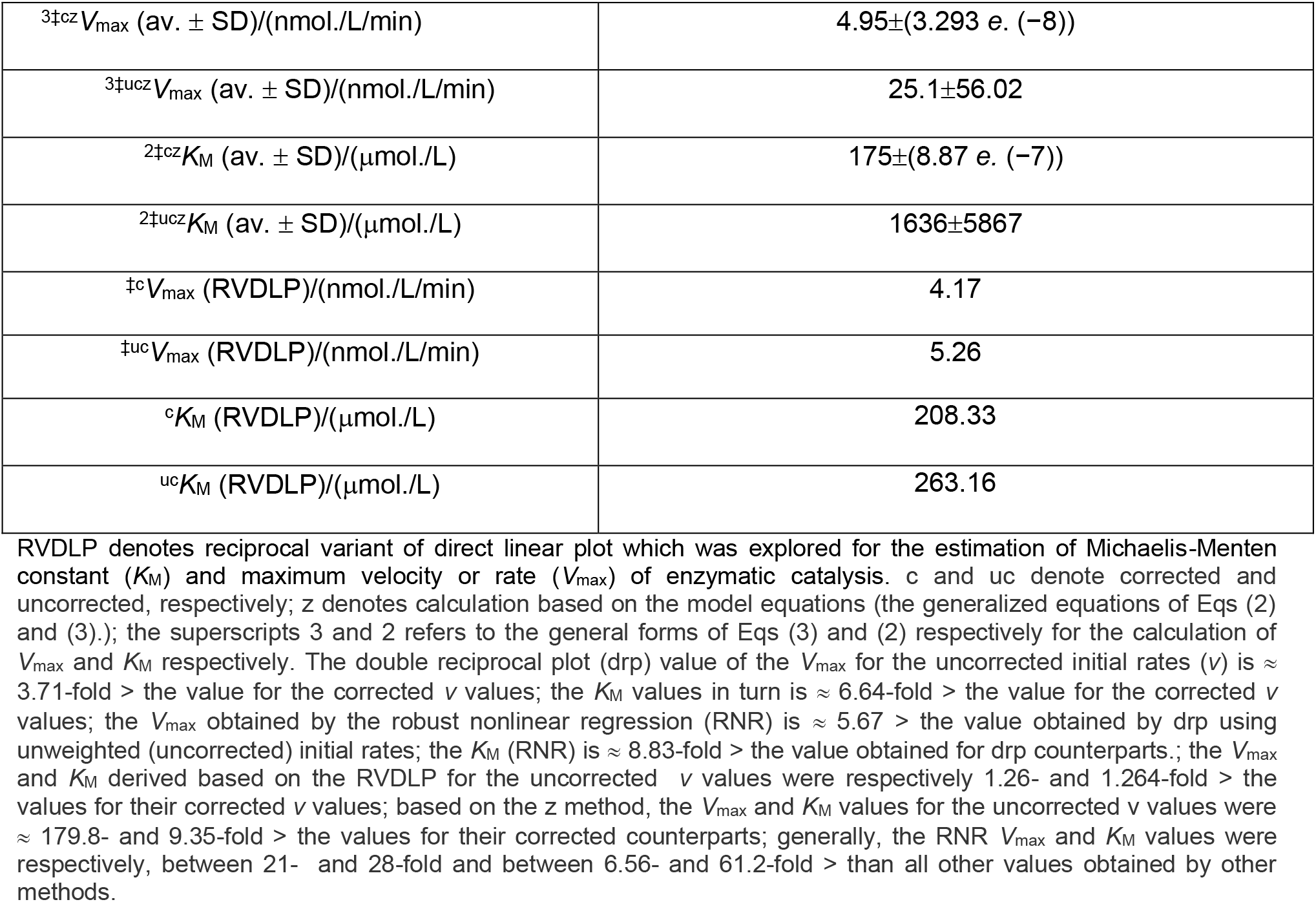
The use of corrected and, uncorrected initial rate data (no replicates) [8] for the determination of kinetic parameters for Tyrosinase by calculation and graphical methods.

Beginning with the double reciprocal plot, the average values of the Michaelian parameters, the *V*_max_ and the *K*_M_, obtained for alpha-amylase by exploring the uncorrected and corrected *v* values are, respectively, 339.6 μmol./L/min and 10.08 g/L for the uncorrected *v* values and 236.7 μmol./L and 5.33 g/L for the corrected *v* values (Table 4). The corresponding median values for three replicates are: 319.66 μmol./L/min and 9.20 g/L for the uncorrected *v* values, and 237.1 μmol./L/min and 5.37 g/L for the corrected *v* values. The calculated values of these kinetic parameters, *V*_max_ and *K*_M_, according to the general equations (general forms of Eqs (2) and (3)) for the uncorrected and corrected *v* values are: 284 μmol./L/min and 7.2 g/L (the corresponding median values are: 291.15 μmol./L/min and 6.13 g/L) and 236.43 μmol./L/min and 5.38 g/L (the corresponding median values are: 234.46 μmol./L/min and 5.22 g/L), respectively The values of the same parameters obtained by RVDLP for the uncorrected and corrected *v* values are: 320.3 μmol./L/min and 8.87 g/L (the corresponding median values are: 3.33 μmol./L/min and 7.94 g/L) and 2.33 μmol./L/min and 5.18 g/L (the corresponding median values are: ≈233 μmol./L/min and 5.14 g/L), respectively. It can be seen that all average and median values obtained from exploring the corrected *v* values are less than those obtained from exploring the uncorrected *v* values.

**Table 4:** The use of corrected and, uncorrected initial rate data (in triplicates) for the determination of kinetic parameters for alpha-amylase by calculation and graphical methods in this study.

|  |  |
| --- | --- |
| $^{cdrp}V_{\max}(\text{av.} \pm \text{SD})/(\mu\text{mol./L/min})$ | $2.367 \pm 0.128(\text{median value} = 0.128)$ |
| $^{ucdrp}V_{\max}(\text{av.} \pm \text{SD})/(\mu\text{mol./L/min})$ | $3.396 \pm 0.292(\text{median value} = 3.197)$ |
| $^{cdrp}K_M(\text{av.} \pm \text{SD})/(\text{g/L})$ | $5.326 \pm 0.763(\text{median value} = 5.374)$ |
| $^{ucdrp}K_M(\text{av.} \pm \text{SD})/(\text{g/L})$ | $10.075 \pm 1.504(\text{median value} = 9.203)$ |
| $^{3cz}V_{\max}(\text{av.} \pm \text{SD})/(\text{nmol./L/min})$ | $236.433 \pm 39.478(\text{median value} = 234.46)$ |
| $^{3ucz}V_{\max}(\text{av.} \pm \text{SD})/(\mu\text{mol./L/min})$ | $284.568 \pm 36.286(\text{median value} = 291.15)$ |
| $^{2cz}K_M(\text{av.} \pm \text{SD})/(\text{g/L})$ | $5.377 \pm 0.228(\text{median value} = 5.224)$ |
| $^{2ucz}K_M(\text{av.} \pm \text{SD})/(\text{g/L})$ | $7.199 \pm 1.6122(\text{median value} = 6.130)$ |
| $^cV_{\max}(\text{RVDLP})/(\mu\text{mol./L/min})$ | $233.02 \pm 11.33(\text{median value} = 2.326)$ |
| $^{uc}V_{\max}(\text{RVDLP})/(\mu\text{mol./L/min})$ | $320.25 \pm 18.48(\text{median value} = 2.371)$ |
| $^cK_M(\text{RVDLP})/(\text{g/L})$ | $5.175 \pm 0.567(\text{median value} = 5.135)$ |
| $^{uc}K_M(\text{RVDLP})/(\text{g/L})$ | $8.869 \pm 0.656(\text{median value} = 9.333)$ |

RVDLP denotes reciprocal variant of direct linear plot which was explored for the estimation of Michaelis-Menten constant ( $K_M$ ) and maximum velocity or rate ( $V_{\max}$ ) of enzymatic catalysis. c and uc denote corrected and uncorrected, respectively; z denotes calculation based on the model equations (the generalized equations of Eqs (2) and (3).); the superscripts 3 and 2 refer to the general forms of Eqs (3) and (2) respectively for the calculation of $V_{\max}$ and $K_M$ respectively; drp denotes double reciprocal plot.

## Discussion

General discussion begins with the reminder that, when *v*_n-1_[*S*_T_]_n_ = *v*_n_[*S*_T_]_n–1_, Eq. (14c) cannot be used to calculate the *V*_max_ and *K*_M_. It implies that a plot of *v* versus [*S*_T_] must yield a perfect straight line (a perfect linear regression with a coefficient of determination of 1 (*R^2^* = 1). In such a situation, Michaelian kinetics is not applicable; rather, it can best be described as a pre-steady-state scenario. The focus remains on how accurate (with improved precision) and valid initial rates can be obtained following the assay of an enzyme. As long as the results expected from Eqs (13) and (15c) are not met, any transformation of initial rate data by whatever means cannot be free from error; the effect of error is only minimized by those transformations. Different methods showcase different degrees of precision. It seems difficult to reach any definitive conclusion on the best method out of the two main considerations, that is, the nonlinear regression and direct linear methods.

However, there are two reports in the literature that go to show that any error in the use of nonlinear regression can give overinflated values of the kinetic parameters [8]. This may be as a result of the adoption of the robust nonlinear regression estimator based on the unfamiliar, modified Turkey’s biweight function for determining the parameters of the Michaelian-Menten equation using experimental measurements (initial rates, to be specific) in kinetic assays. Although the procedure for modification is not understood, it is not unlikely that the approach may have overestimated the values of these kinetic parameters considering similar issues in the literature [2], where it is believed that the essence of biweight regression is that after the first iteration, observations with large values of squared residue are assigned decreased weight. If such a procedure is arbitrary, it may be a source of error. It is well known that linear transformation overestimates kinetic parameters; there is convincing evidence in this research and in the literature [2]. In this study, the values of the kinetic parameters obtained from the reciprocal variant (RVDLP) were much less than the values obtained from the double reciprocal plot (drp) but slightly similar to the values obtained using the model equations, that is, the generalized Eqs (2) and (3), both for the corrected and uncorrected initial rates in the literature [8] and in this study. In all cases, the use of uncorrected initial rates gives higher values for the kinetic parameters: For example, when using drp, the *V*_max_ for the uncorrected initial rates (*v*) is ≈ 3.71-fold > the value for the corrected *v* values; the *K*_M_ values in turn is ≈ 6.64-fold > the value for the corrected *v* values; the *V*_max_ obtained by the robust nonlinear regression (RNR) is ≈ 5.67 > the value obtained by drp using unweighted (uncorrected) initial rates; the *K*_M_ (RNR) is ≈ 8.83-fold > the value obtained for drp counterparts; the *V*_max_ and *K*_M_ derived based on the RVDLP for the uncorrected *v* values were respectively 1.26- and 1.264-fold > the values for their corrected *v* values; based on the z-method, the *V*_max_ and *K*_M_ values for the uncorrected *v* values were ≈ 179.8- and 9.35-fold > the values for their corrected counterparts; generally, the RNR *V*_max_ and *K*_M_ values were respectively, between 21- and 28-fold and between 6.56- and 61.2-fold > than all other values obtained by other methods.

The wide disparity observed in the kinetic parameters obtained, based on different methods, for Tyrosinase from *Solanum tuberosum* [8] is totally at variance with the observation with alpha-amylase from *Aspergillus oryzae*. The difference is that a nonlinear approach was not used for the alpha-amylase. The drp values of *V*_max_ and *K*_M_ for the uncorrected *v* values were respectively 1.2- and 1.34-fold > their corrected counterparts; the results from RVDL showed that the *V*_max_ and *K*_M_ for the uncorrected *v* values were respectively 1.3- and 1.714-fold > the values for their corrected counterparts, while the model approach gave values that were 1.2- and 1.34-fold > the values for their corrected counterparts. There are some deductions to be made from this study and the work done using literature [8] information. If the initial rates are accurate, the kinetic parameters generated from linear transformation and RVDLP as well as the z method may be similar; for instance, the values of *V*_max_ for the corrected *v* values are 5.609, 4.95, and 4.17 nmol./L/min (Table 3), which were respectively obtained by the drp, RVDLP, and z methods, are not widely different. Also, the *K*_M_ values, which are 180.93, 175, and 208.6 mmol./L (Table 3), obtained by the drp, RVDLP, and z methods, are not widely different. Pieces of information in the literature support this observation. In one case, the kinetic parameters obtained from drp were > values from other methods (from the study of nutrient uptake by algal cells), while in another case, values from drp were < values from other methods, and in both cases, known models were fitted to initial rates from the papain assay [3].

Worries about outliers and the common practice of basing weighting schemes on hypotheses about how error variances vary with true rates, which may be incorrect because of variations in experimental conditions, technique, and skills [2], should be things of the past in the light of this new model. Statistical analysis based on Gaussian statistics requires a large number of replicates; 6 to 10 assays per concentration of the substrate can be very tedious if manually conducted; a lot of time and resources, to be specific, can translate into a huge cost. This has been the concern of some researchers [19], who posit that the implications of computer control can promote rapid characterization throughputs and cost savings; the use of automation can reduce or eliminate errors, and as such, give a more reliable result. In order to be more compliant with Gaussian statistics, both speed and precision can be achieved with a single assay per substrate concentration as long as the substrate concentration range covers more than eight different concentrations. According to the new models, the effective population size is “*n*-1” where *n* is the number of different concentrations of the substrate. However, while computer-aided assay provides accurate initial rates, it is also prudent to evaluate the data obtained by fitting Eqs (13) and (14c) to the initial rates to ensure that *v*_n–1_[*S*_T_]_n_ is not equal to *v*_n_ [*S*_T_]_n–1_ (Eq. (14c)) and that decreasing trends are observed as expressed by Eq. (13). The same concern is applicable to Eq. (15 c). The concern about Eq. (9) and (12) is that they are quite complex and should take a longer computational time; otherwise, both can always reproduce the parameter they stand for if the initial rates are very accurate, as applicable to Eq. (2) and (3). The application of the pseudo-statistical approach represented by Eqs (40) and (41) is reserved for a future investigation.

## Conclusion

In this study, the equations for the correction of initial rates where necessary, the equations that can aid the identification of error(s) in the initial rates, alternative equations based on the Michaelian equation, and “pseudo-statistical” equations were derived. The equations that were evaluated were found to be valid in that the corrected initial rates for which errors were identified were able to give kinetic parameters similar to the reference standard values generated from fitting the reciprocal variant of a direct linear plot to the corrected initial rates. When fitted to the correct initial rates, the double reciprocal and the new Michaelis-Menten based derived equation (or model) can produce comparable results if the initial rates were correctly determined. It is recommended to use a broad substrate concentration range of 6 to “*n*” (where *n* ⩾ 10). A single enzyme assay can be conducted for each substrate concentration without violating the requirement for Gaussian statistics. This method saves both time and money. A future study will attempt to evaluate, among other questions, the “pseudo-statistical” equations.

## Materials and Methods

### Materials

#### Chemicals

The enzyme which was assayed is *Aspergillus oryzae* alpha-amylase (EC 3.2.1.1) and, insoluble potato starch, was the substrate; both were purchased from Sigma–Aldrich, USA. Tris 3, 5—di-nitro-salicylic acid, maltose, and sodium potassium tartrate tetrahydrate were purchased from Kem Light Laboratories in Mumbai, India. Hydrochloric acid, sodium hydroxide, and sodium chloride were purchased from BDH Chemical Ltd., Poole, England. Distilled water was purchased from the local market. Distilled water was purchased from the local market. As a word of caution, readers of this paper should be aware that the use of the same enzyme in articles by the same author(s) is strictly due to budgetary constraints; however, this is not a serious concern because each paper addresses different issues, such as the evaluation of new models.

#### Equipment

An electronic weighing machine was purchased from Wensar Weighing Scale Limited, and a 721/722 visible spectrophotometer was purchased from Spectrum Instruments, China; a *p*H meter was purchased from Hanna Instruments, Italy.

### Methods

#### Preparation of reagents and assay

The method of assaying the enzyme is Benfield’s method [20] Gelatinized potato starch, whose concentration range is 5–10 g/L was used as substrate. The reducing sugar produced upon hydrolysis of the substrate at 22 °C using maltose as a standard was determined at 540 nm with an extinction coefficient approximately equal to 181 L/mol.cm. The assay took 3 minutes to complete. A mass concentration equal to 2.5 mg/L of *Aspergillus oryzae* alpha-amylase was prepared in a Tris-HCl buffer at *p*H 6.9; there were no special considerations in the choice of *p*H and temperature. As in previous publications [16, 17], the evaluation of new equations was the only overriding interest. In this regard, alpha-amylase was assayed in this study while initial rate values for Tyrosinase from *Solanum tuberosum* were as reported in the literature [8]. Lineweaver-Burk [21] and the reciprocal variant of direct linear plot [7] were explored for the generation of kinetic parameters.

## Abbreviations

err: error
*e*: exponent
drp: double reciprocal plot
RVDLP: reciprocal variant of direct linear plot
RNR: robust nonlinear regression
z: denotes calculation based on the model equations
*v*: initial rate
*K_M_*: Michaelis-Menten constant
*V*_MAX_: maximum rate of catalysis
c: corrected
uc: uncorrected
QSSA: quasi-steady-state assumption
rQSSA: reverse quasi-steady-state assumption
sQSSA: standard quasi-steady-state assumption
L-DOPA: L-3, 4-dihydroxyphenyl alanine
*e*_i_: random error component of initial rates.

## Author contribution

The sole author designed, analyzed, interpreted and prepared the manuscript.

## Acknowledgments

The management of the Royal Court Yard Hotel, Agbor, Delta State, Nigeria, is immensely appreciated for the supply of electricity during the preparation of the manuscript. The provider of the QuillBot grammar checker is thanked for the proofreading services.

